# Virus propagation linked to exceedingly rare gene-expression errors

**DOI:** 10.1101/2024.05.21.595180

**Authors:** Raquel Luzón-Hidalgo, Gianluca d’Agostino, Valeria A. Risso, Asuncion Delgado, Beatriz Ibarra-Molero, Luis A. Campos, Jose Requejo-Isidro, Jose M. Sanchez-Ruiz

**Affiliations:** Departamento de Quimica Fisica. Facultad de Ciencias, Unidad de Excelencia de Quimica Aplicada a Biomedicina y Medioambiente (UEQ), Universidad de Granada 18071 Granada, Spain; Centro Nacional de Biotecnología (CNB), CSIC, 28049 Madrid, Spain; IMDEA-Nanociencia, Ciudad Universitaria de Cantoblanco, 28049 Madrid, Spain; Unidad de Nanobiotecnología, CNB-CSIC-IMDEA Nanociencia Associated Unit, 28049 Madrid, Spain

## Abstract

Viruses are obligate parasites that establish extensive interactions with proteins and other biomolecules of their hosts. About 20% of protein molecules bear phenotypic mutations due to errors during gene expression. Phenotypic mutations are not inherited and are not purged/amplified by natural selection. Therefore, protein variants harboring phenotypic mutations remain at very low levels. Here, we show that proteins at exceedingly low levels may enable virus propagation. Bacteriophage T7 recruits the host thioredoxin as an essential processivity factor for its replisome. Thioredoxin constitutive expression yields 10000-20000 molecules per *E. coli* cell. We inserted early stop codons in the thioredoxin gene and appended to its end the sequence encoding for a photoconvertible fluorescent protein. Virus propagation was not abolished, indicating that some thioredoxin molecules were produced through mistranscription or mistranslation. Single-molecule localization microscopy detected 12±5 molecules per cell when an ochre codon was inserted. This work demonstrates that crucial virus-host biomolecular interactions may need occur only a few times to trigger virus propagation and supports that viruses may exploit the wide diversity of host and viral protein variants arising from gene-expression errors to establish such interactions. Immediate implications of this notion for the mechanisms of cross-species transmission and antibody evasion are discussed.

## Main

Bacteriophage T7 is a lytic phage that infects strains of *E. coli* and destroys the infected cells, with the concomitant release of more than a hundred of new virions per infected cell (Quimron et al., 2010). Infection starts with the attachment of a virion to the cell surface followed by transfer of the viral DNA to the cytosol. This triggers processes that eventually lead to the death of the cell. In addition to infection, phage T7 propagation requires, among other processes, the assembly inside the host cell of a viral replication machinery, in such a way that copies of the viral DNA are generated to be subsequently packed inside the new virions. The minimalist replisome of bacteriophage T7 consists of a DNA polymerase, a helicase/primase, a ssDNA-binding protein and a thioredoxin (Hamdan and Richardson, 2009). Thioredoxins are small redox proteins involved in a diversity of processes in all known cells (Holmgren, 1985; Kumar and Richardson, 2004). Crucially, the thioredoxin in the replisome of phage T7 is a protein from the *E. coli* host that is recruited by the virus to serve as an essential processivity factor for the viral DNA polymerase (Fig. 1a) (Etson et al., 2010). *E. coli* thioredoxin (Fig. 1b) is expressed in *E. coli* to copy numbers of about 10000-20000 molecules per cell (Holmgren, 1981; Lunn et al., 1984). The immediate question we asked in this work is whether replisome assembly and subsequent replication of phage T7 requires the presence of ∼10^4^ thioredoxin molecules in a host cell or whether a much smaller number of thioredoxin molecules per cell suffices to trigger virus propagation. The answer we obtained to this question has general implications for our understanding of the mechanisms of virus adaptation.

**Fig. 1.**
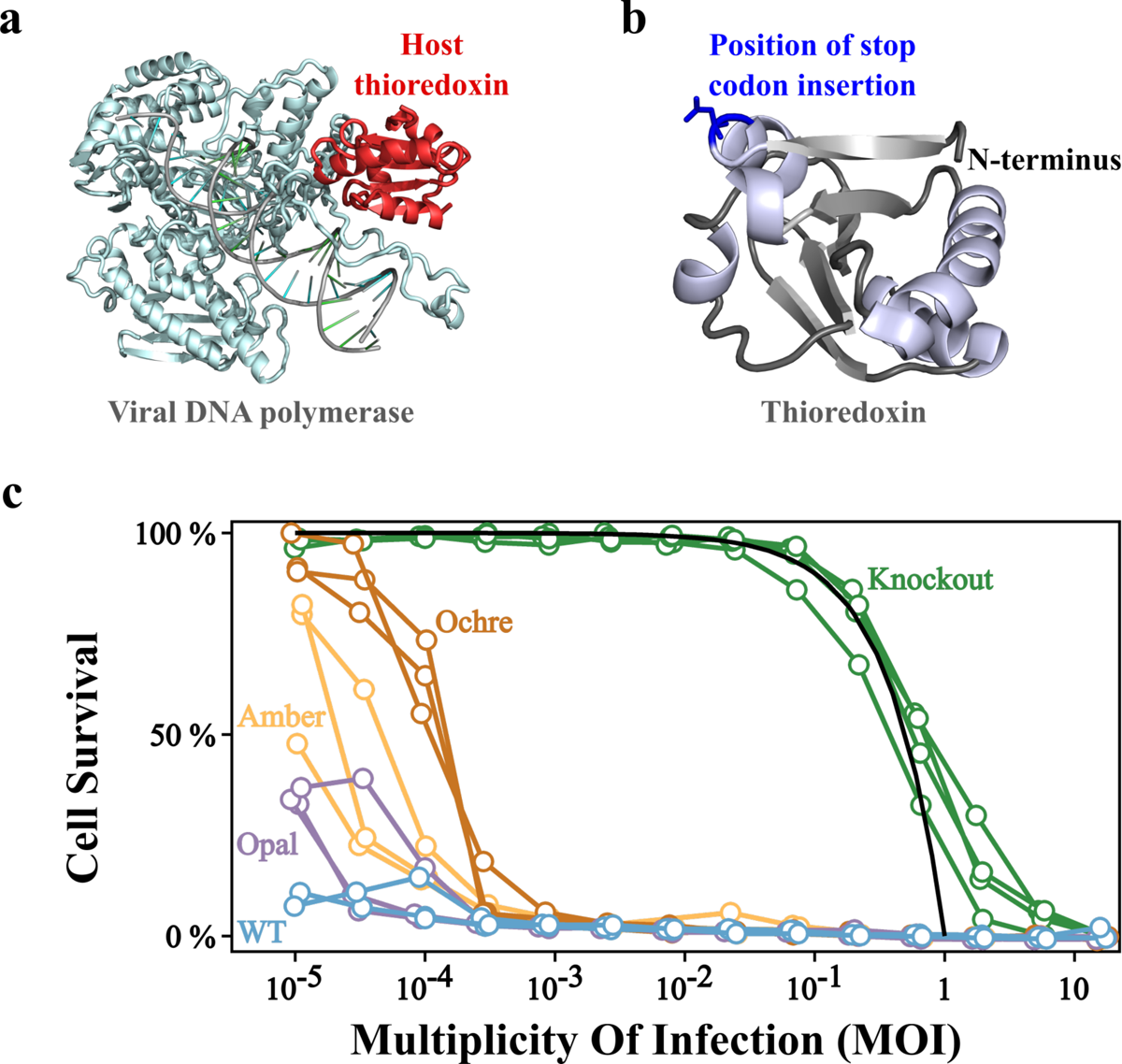
Host-factor engineering of phage T7 propagation. **a**, Structure of the viral DNA polymerase of phage T7 interacting with the host thioredoxin (PDB ID 1T8E). Thioredoxin functions as an essential processivity factor for the polymerase (Etson et al., 2010). The interaction of the two proteins is tight with a reported dissociation constant of 5 nanomolar (Hamdan and Richardson, 2009). **b,** 3D-structure of *E. coli* thioredoxin (PDB ID 2TRX) showing the position of insertion of stop codon stop (position 11). **c,** Propagation of phage T7 in *E. coli* cells with modified expression of the host factor. Experiments were performed with knockout Trx^-^ cells, in which thioredoxin is absent, with cells in which the thioredoxin gene had been modified through the insertion of opal, amber and ochre stop codons, and with cells transformed with the unmodified (wt) thioredoxin gene. For each type of cell, propagation experiments with different initial multiplicity of infection (MOI) values were set up and the extent of cellular death was assessed by turbidimetry after 1 hour (see Methods for details). For each type of cell, three propagation profiles (fraction of cell survival vs. MOI) were determined using three biological replicates (4 biological replicates for the knockout cells). Phage does not replicate in knockout Trx^-^ cells, since they do not express the essential host factor, thioredoxin. Consequently, death of knockout cells requires MOI values approaching unity. In fact, the continuous line is the theoretical prediction if the fraction of cell death equals the MOI value. By contrast, complete death is observed for all the other types of cells, even with very low MOI values. An efficiency of propagation ranking of ochre<amber<opal<wt is visually apparent and likely reflects the generated thioredoxin levels.

To address the question posed above, we made use of a previously characterized (Delgado et al., 2017) knockout *E. coli* strain in which thioredoxin genes have been deleted. We shall refer to this strain as *E. coli* Trx^-^. This strain can grow, albeit more slowly than wild-type *E. coli*, likely because the glutaredoxin pathway substitutes to some extent the functions of thioredoxin. *E. coli* Trx^-^ effectively decouples virus infection from virus replication. That is, *E. coli* Trx^-^ cells are infected by the phage and die as a result of the infection, but they cannot amplify the virus, since the absence of thioredoxin precludes the assembly of efficient viral replisomes.

Consequently, death of all the cells in a sample of *E. coli* Trx^-^ requires a large multiplicity of infection (MOI: the ratio of number of virus particles to number of host cells) of about unity or larger, as it is visually apparent in Fig. 1c. By contrast, all cells in a wild-type *E. coli* sample are eventually killed by the phage, even in experiments with very low MOI values, simply because the synthesis of thioredoxin molecules enables replisome assembly and, consequently, initial infection of just a few cells generates many new virions that propagate the infection.

We complemented the knockout *E. coli* Trx^-^ cells with a plasmid harboring the gene of *E. coli* thioredoxin, which allows us to easily modify thioredoxin in several useful ways. The basal expression of the pET plasmid we used already produces thioredoxin levels similar to those corresponding to the constitutive expression in wild-type *E. coli* cells (see Delgado et al., 2017 and results given further below). In fact, even under non-inducing conditions, complementation restores the phage infection pattern of wild-type *E. coli*, *i.e.,* essentially all cells eventually die in experiments starting with very low MOIs (Fig. 1c). In view of this result, we decided to decrease the expression levels by introducing early stop codons in the thioredoxin gene. It has been known for many years that stop codons are “leaky” to different extents as a result of, for instance, transcription errors or translational readthrough (Parker, 1989). Therefore, we hypothesized that early stop codons would not completely block thioredoxin synthesis but would lead to substantially decreased thioredoxin copy numbers.

Position 11 was selected for stop-codon insertion because it is exposed to the solvent in the 3D-structure of thioredoxin (Fig. 1b) and shows considerable diversity among thioredoxins from different species (Extended Data Fig. 1). Therefore, different amino acid residues at position 11 (result of the misreading of the stop-codon) are likely to lead to correctly folded thioredoxin molecules. We introduced opal, amber and ochre stop codons at position 11 and found clear evidence of virus propagation in the three cases (Fig. 1c and Extended Data Fig. 2). The number of plaque-forming units (PFU) for Trx^-^ cells transformed with opal-trx, amber-trx and ochre-trx genes were thus on the same order as the PFU values for cells transformed with wild type thioredoxin. However, propagation was less efficient when stop codons had been inserted.

Furthermore, among the genes with stop codons inserted, propagation was least efficient with the ochre stop codon and most efficient with the opal stop codon. This is clearly shown by the profiles of cell survival versus MOI (Fig. 1c) and by the size of the plaques (Extended Data Fig. 2) in experiments aimed at determining the number of PFU values. Remarkably, the observed ranking of propagation efficiency ochre<amber<opal (Fig. 1c and Extended Data Fig. 2) matches the known ranking of stop-codon leakiness in *E. coli*. That is, opal is known to be particularly leaky while ochre shows the lower readthrough frequency (Parker, 1998). The congruence supports that the differences in propagation efficiency reflect the number of generated thioredoxin molecules and consequently the number of assembled replisomes.

In a first attempt to quantify the number of thioredoxin molecules enabling virus propagation we appended to the end of the thioredoxin gene (Fig. 2a) the sequence encoding for the enhanced green fluorescent protein (eGFP). In this way, every synthesized thioredoxin molecule should carry an attached eGFP molecule and eGFP fluorescence could provide a metric of the amount of thioredoxin generated. As expected, knockout Trx^-^ cells transformed with wt-trx-eGFP show high fluorescence levels (Fig. 2a). On the other hand, introduction of an ochre stop at position 11 of thioredoxin (Fig. 2b) leads to fluorescence levels that are very low and similar, for almost all cells, to those observed with the knockout *E. coli* Trx^-^ cells (Fig. 2c). This indicates that the contributions from endogenous autofluorescence and eGFP are comparable and that only a small fraction of ochre-trx-eGFP transformed cells expressed eGFP to a detectable level in these experiments (Extended Data Fig. 3).

**Fig. 2.**
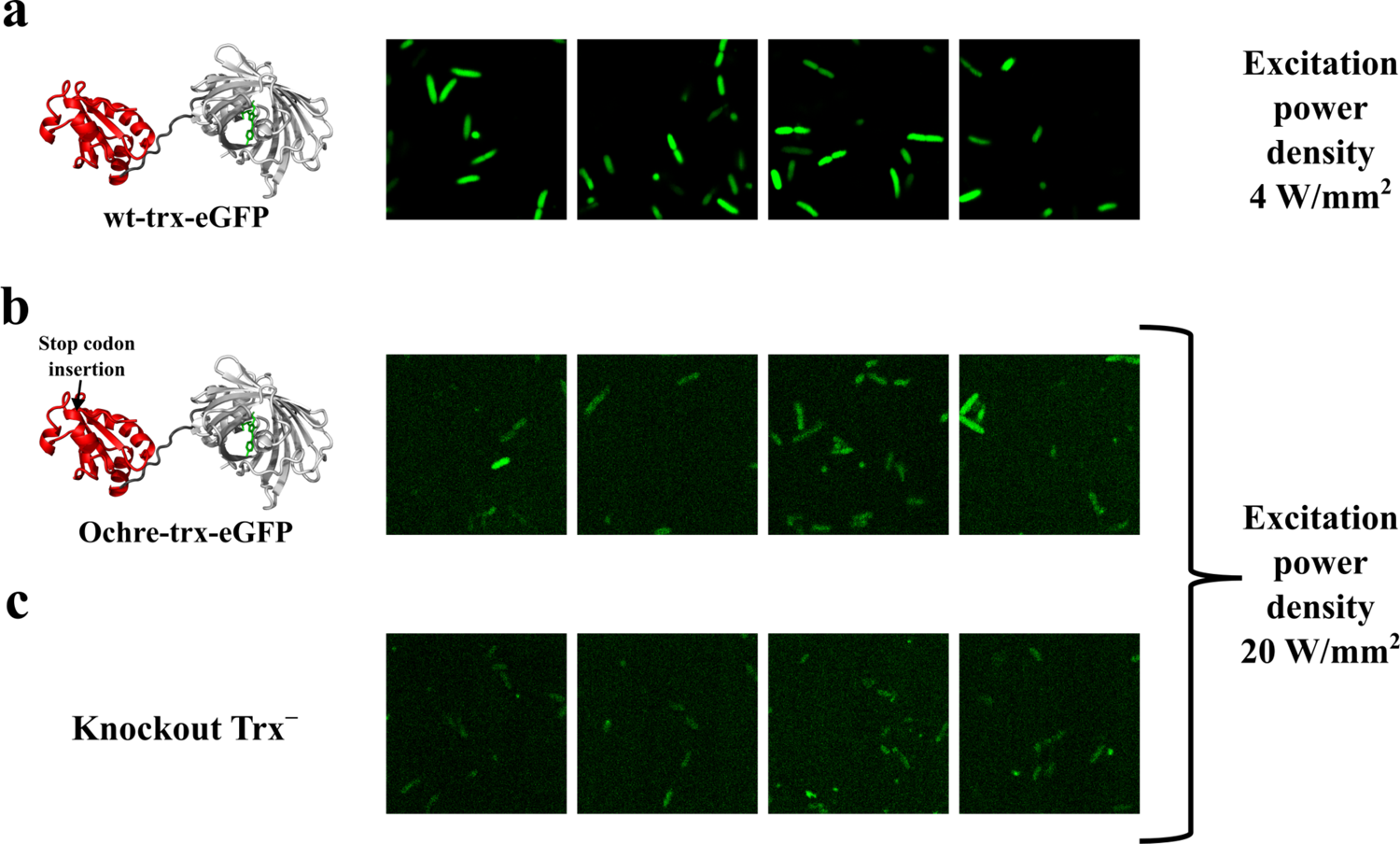
Assessment of thioredoxin levels using eGFP fusions. **a**, Depiction of the 3D-structure of the wt-trx-eGFP fusion construct and representative microscopy images of cells transformed with wt-trx-eGFP. **b,** Depiction of the 3D-structure of the ochre-trx-eGFP fusion construct with the position of insertion of the ochre codon highlighted and representative microscopy image of cells transformed with ochre-trx-eGFP. **c,** representative microscopy images of knockout Trx^-^ cells. See Extended Data Fig. 3 for further details.

While it is clear from these experiments that the level of trx-eGFP constructs in cells transformed with ochre-trx-eGFP is indeed very low, it does not appear possible to use eGFP fluorescence to estimate the number of generated molecules per cell.

In view of the results described in the previous paragraph, we decided to append to the thioredoxin gene the sequence encoding for the photoconvertible fluorescent protein mEos2 (McKinney et al., 2009). In this way the synthesized thioredoxin molecules are linked to this protein (Fig. 3a), as we ascertained from the UV-VIS spectra of the purified fusion construct (Fig. 3a). Crucially, the presence of mEos2 in the final protein product of thioredoxin gene expression allows us to use single-molecule localization microscopy (SMLM) (Lelek et al., 2021) to quantify the number of generated thioredoxin molecules. We first applied the approach to cells expressing wild type thioredoxin molecules (*i.e.,* no stop codon at position 11). That is, we transformed the knockout Trx^-^ cells with the wt-trx-mEos2 gene. The presence of the attached mEos2 does not preclude the death of all cells in virus propagation experiments starting with low MOI values down to ∼10^-4^ (Fig. 3b; see also Extended Data Fig. 2). To quantify the number of thioredoxin molecules generated (strictly, the number of wt-trx-mEos2 fusion constructs), we simultaneously photoconverted mEos2 in fixed cells using 405 nm constant illumination and imaged the photoconverted mEos2 at 561 nm using Total Internal Reflection (TIRF) configuration until all mEos2 molecules within the TIRF volume were bleached (Supplementary Video 1).

**Fig. 3.**
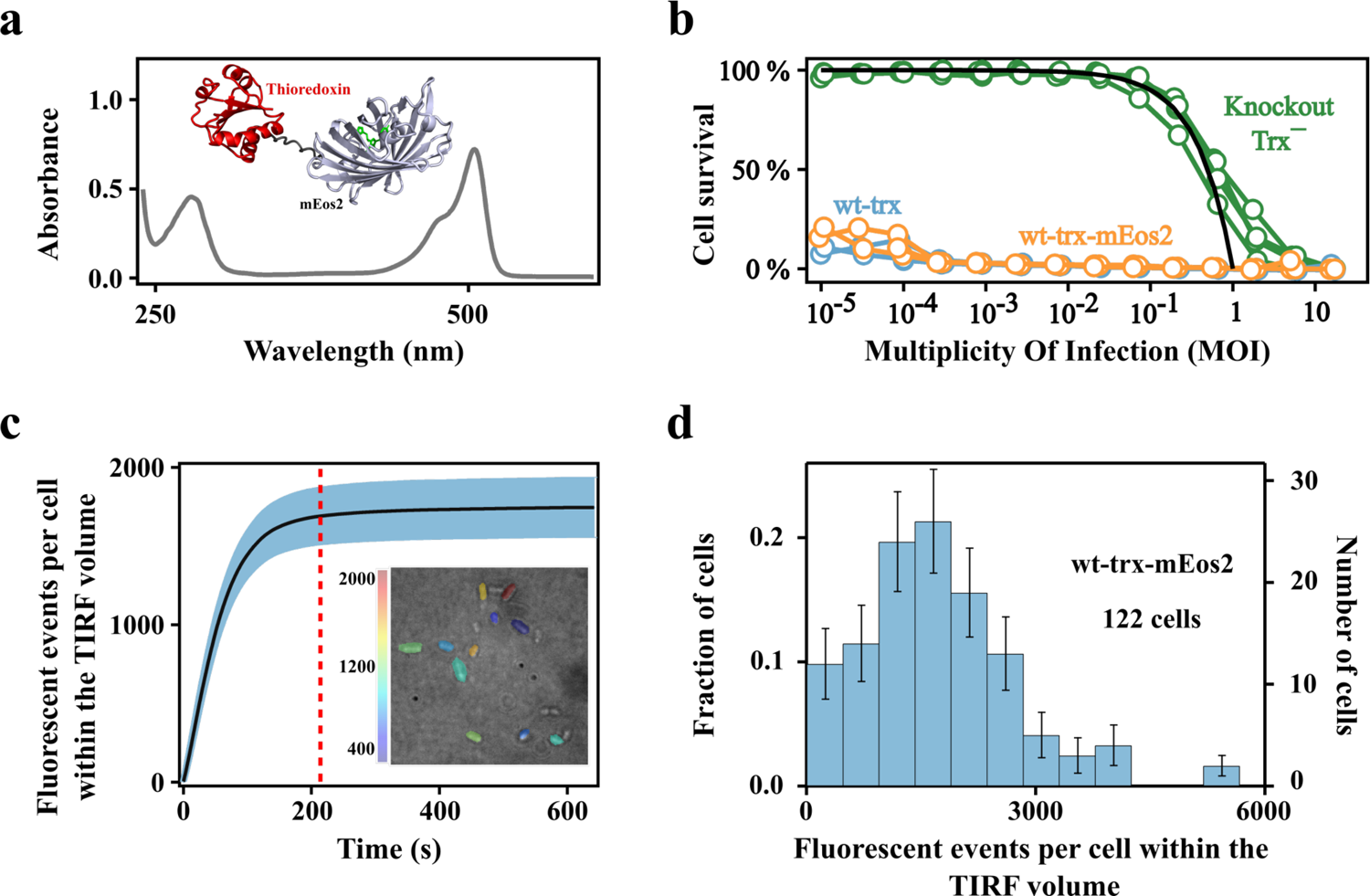
Single-molecule localization in cells transformed with wt-trx-mEos2. **a**, Representative UV-VIS spectrum and depiction of the 3D-structure of the wt-trx-mEos2 fusion construct. The construct was prepared following a cell growth protocol similar to that used for virus propagation and microscopy experiments. Three independent preparations led to a 506 nm to 280 nm absorbance ratio 1.45±0.30, which is consistent with a 1:1 mEos2:thioredoxin stoichiometry (see Methods for details). **b,** Propagation of phage T7 in cells transformed with wt-trx and wt-trx-mEos2. These experiments follow the same protocol as those of Fig. 1c and show that the attached mEos2 does not preclude the death of all cells in virus experiments starting with low MOI values. **c,** Cumulative number of fluorescent events per cell within the TIRF illumination volume derived from localization microscopy experiments with cells transformed with wt-trx-mEos2. The shadowed area is the 95% confidence interval, calculated by data bootstrapping. The dashed line indicates the time for quantification of fluorescent events at the plateau (t=214s). The inset is a representative image with individual cells colored according to the number of fluorescent events detected. **d,** Distribution of fluorescent events per cell within the TIRF volume. Data on 122 cells were used to derive the distribution. For representation purposes, the experimental range of fluorescent events has been divided into 12 bins. The uncertainties bars at each bin are based on counting statistics.

Independent FRAP (fluorescence recovery after photobleaching) experiments (Extended Data Fig. 4) showed that fluorescent proteins did not diffuse in cells during the experiments. Consequently, only the molecules present within the TIRF illumination depth are detected. Since this depth was set to 150 nm, *i.e.*, 6.5 times smaller than the relevant length of the *E. coli* cell (Extended Data Fig. 5 and Methods), the number of trx-Eos2 copies measured in TIRF configuration had to be scaled by that factor to estimate the total number of molecules present in the cell.

The cumulative count of fluorescent events in cells transformed with wt-trx-mEos2 quickly grew before reaching a plateau (Fig. 3c), indicating that there were no remaining fluorescent molecules. We thus quantified the number of fluorescent events in every cell at the plateau (Fig. 3c-3d). Fig. 3d summarizes the results of the determination of the number of thioredoxin molecules (strictly, the number of fluorescent thioredoxin-mEos2 fusion constructs) for 122 cells. The wide distribution observed reflects the fact that gene expression is an essentially stochastic phenomenon (Cai et al., 2006), although differences in the number plasmid copies per cell may also contribute. Most cells displayed substantial levels of thioredoxin expression, with an average value of 1737±198 (95% confidence interval, CI) molecules per cell within the TIRF volume. Considering the relation between the TIRF volume and the size of an *E. coli* cell (Extended Data Fig. 5 and Methods), this amounts to roughly 11000 molecules per cell, a value on the order of the constitutive expression levels for thioredoxin in wild-type *E. coli*: ∼10000-20000 molecules per cell (Holmgren, 1981; Lunn et al., 1984).

We now appended the mEos2 sequence to the ochre-containing thioredoxin gene (Fig. 4a) and set out to quantify thioredoxin expression as we had previously done when no stop codon was present. That is, we transformed the knockout Trx^-^ cells with the ochre-trx-mEos2 gene. The attachment of mEos2 is not expected to change the number of generated thioredoxin molecules but only to provide a way to quantify them. This purpose would be served even if, due perhaps to steric interference, the presence of mEos2 precluded the interaction of thioredoxin with the viral DNA polymerase. In reality, we observed that cells transformed with ochre-trx-mEos2 could still sustain virus propagation, although less efficiently than cells transformed with ochre-trx (Fig. 4b and Extended Data Fig. 2). In contrast to the determination of wt-trx-mEos2 copy number with no stop codon (Fig. 3), the number of fluorescent events registered in cells expressing the ochre-trx-mEos2 construct was extremely low and of the same order as the number of fluorescent events observed in knockout Trx^-^ cells (Fig. 4c and Supplementary Videos 2 and 3). In addition, the cumulative count of fluorescence events in both cases, cells transformed with ochre-trx-mEos2 and knockout cells did not reach a plateau (Fig. 5a). On the contrary, after the initial asymptotic exponential increase phase, the cumulative count kept increasing in a linear fashion. This linear contribution was almost undetectable in cells expressing wt-trx-mEos2 because of the much higher number of events associated to the asymptotic exponential phase (Fig. 3c).

**Fig. 4.**
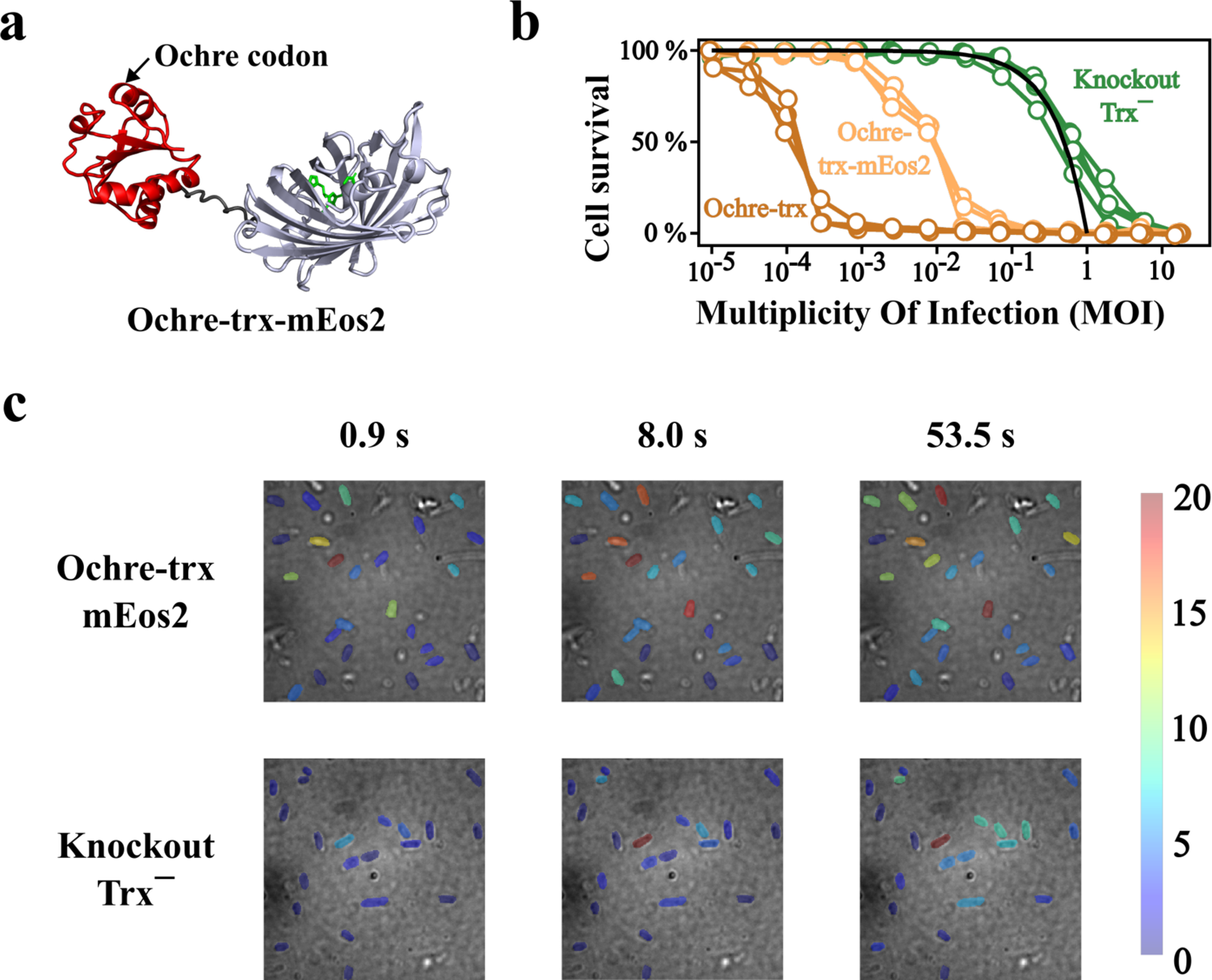
Single-molecule localization in cells transformed with ochre-trx-mEos2. **a**, Depiction of the 3D-structure of the ochre-trx-mEos2 construct showing the position of insertion of the ochre codon. **b,** Propagation of phage T7 in cells transformed with ochre-trx and with ochre-trx-mEos2. These experiments follow the same protocol as those of Fig. 1c and show that, while the attached mEos2 impairs to some extent the cells capability to sustain virus propagation, it does not preclude complete cell death in experiments starting with low MOI values. **c,** Representative images obtained in localization microscopy experiments involving cells transformed with ochre-trx-mEos2 and knockout Trx^-^ cells at increasing times during the photoactivation and acquisition sequence. Individual cells are colored according to the number of fluorescent events detected. These images are shown for illustration only. A quantitative analysis of the data is given in Fig. 5.

**Fig. 5.**
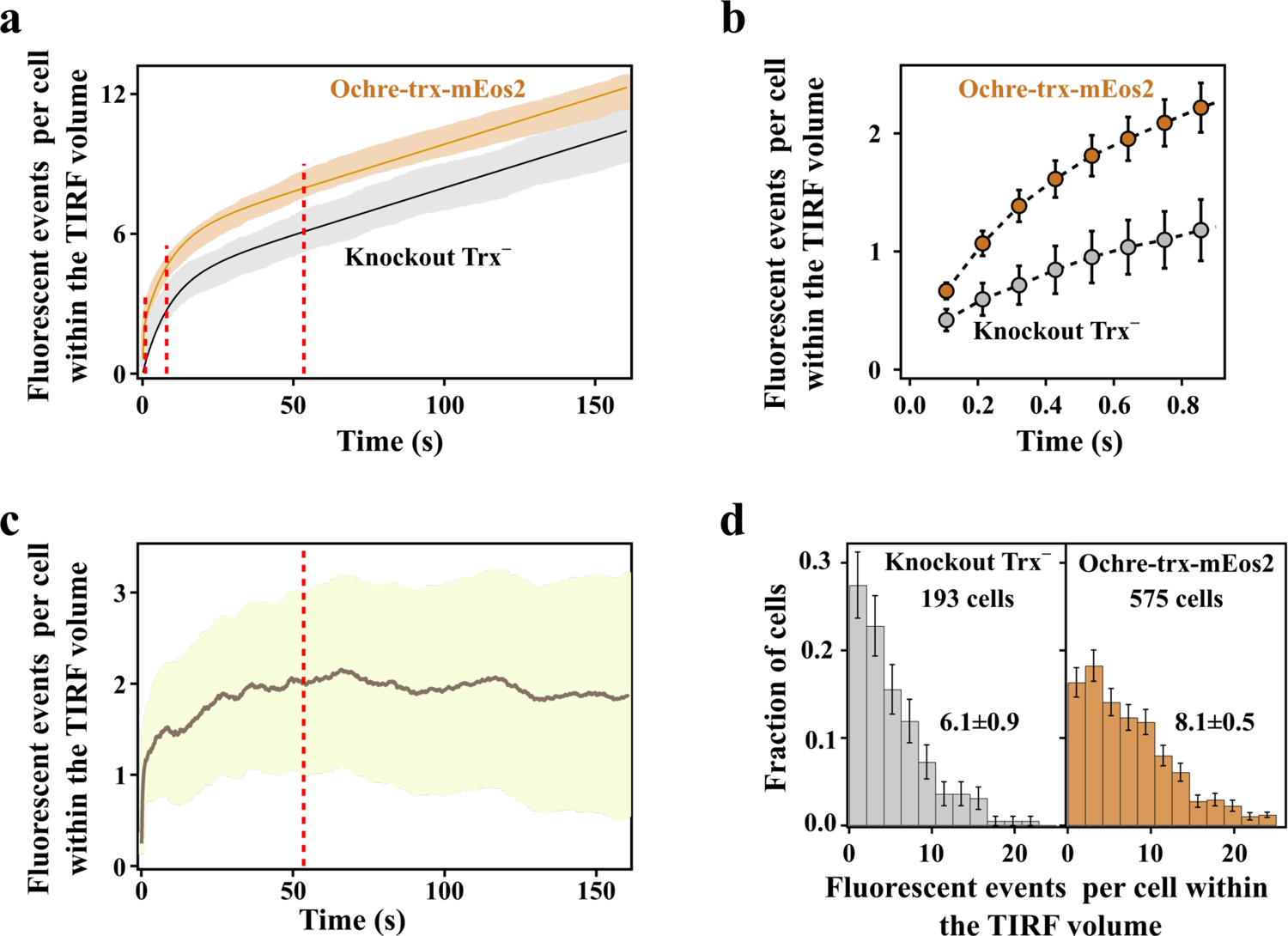
Quantification of thioredoxin molecules in cells transformed with ochre-trx-mEos2. **a**, Number of fluorescent events per cell within the TIRF volume derived from photobleaching experiments with knockout Trx^-^ cells and with cells transformed with ochre-trx-mEos2. For knockout Trx^-^, data for 193 cells were obtained in 3 independent experiments involving 3 biological replicas. For ochre-mEos2, data for 575 cells were obtained in 5 independent experiments involving 5 biological replicas. The shadowed regions show the 95% confidence interval of the experimental data (calculated by bootstrappoing). The solid lines are the cumulative exponential-linear fit (equations 1 and 2) fitted globally to the ochre-wt-mEos2 and knockout cells counts. **b,** Blowup of the low-time region of the plot in a, highlighting the much faster initial increase of cumulative number of fluorescent events for the cells transformed with ochre-trx-mEos2 as compared with the knockouts. **c,** Difference between the profiles for ochre-trx-mEos2 and knockout Trx^-^ shown in panel a. The solid line is the average value and shadowed area is the 95% confidence interval. The profile stabilizes at 2±1 (95% CI) fluorescence events per cell within the TIRF volume. **d,** Distribution of fluorescent events per cell within the TIRF volume for knockout Trx^-^ cells and for cells transformed with ochre-trx-mEos2. The distributions shown correspond to a time of 53.5 s. For representation purposes, the experimental range of fluorescent events has been divided into 12 bins. The uncertainties bars at each bin are based on counting statistics.

There are, however, two clear differences between the profiles of cumulative fluorescent events for cells transformed with ochre-trx-mEos2 and the knockout cells. First, the number of fluorescent events per cell within the TIRF volume is consistently higher for cells expressing ochre-trx-mEos2, as is apparent in Fig. 5a. Second, the number of fluorescent events initially increases at a much faster rate for cells expressing ochre-trx-mEos2 as compared with the knockouts (Fig. 5b). It is important to note here that, for cells expressing ochre-trx-mEos2, we performed 5 independent experiments with 5 biological replicates, analyzing around 100 cells in each experiment (575 total). Likewise, for knockout Trx^-^ cells, we performed 3 independent experiments with 3 biological replicates, probing around 60 cells in each experiment (193 total). The agreement among the different experiments performed with each type of cells was excellent, as shown by the 95% CI in Figs. 5a and 5b. Therefore, the differences we have noted above (higher number of fluorescent events and higher initial rates for cells expressing ochre-trx-mEos2 as compared with knockouts) are quite robust results.

Overall, our results (Figs. 4 and 5) are consistent with a significant contribution of endogenous autofluorescence, scattering and even detector noise to the cumulative fluorescence events for both, cells transformed with ochre-trx-mEos2 and knockout Trx^-^ cells. They also reveal, however, an additional contribution in the former case due to exogenous fluorescence linked to the trx-mEos2 molecules generated through mistranscription or mistranslation of the ochre codon. To evaluate this contribution, we performed a global least-squares fitting of the two profiles based on the following equations for asymptotic exponential growth with a linear slope,

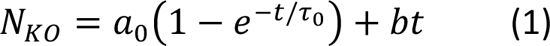

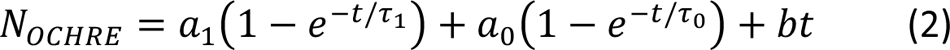

where N_KO_ and N_OCHRE_ are the cumulative numbers of fluorescent events for knockout Trx^-^ cells and cells transformed with ochre-trx-mEos2, respectively, and t is time. The terms *a*_0_(1 − *e*^t/τ_0_^) and *bt* represent contributions from processes other than the photoconversion of trx-mEos2 molecules and are common to the two equations. The term *a*_1_(1 − *e*^t/τ_1_^) is the contribution of trx-mEos2 photoconversion and appears only in the equation for N_OCHRE_. The two equations were simultaneously fitted to the profiles of N_KO_ and N_OCHRE_ imposing that the common parameters (a_0_, ι−_0_ and b) must have the same values for the two profiles. The fit was visually excellent (Fig. 5a) and led to the following values for the parameters describing the photoconversion term: a_1_=1.9±0.7 (95% CI) fluorescent events per cell within the TIRF volume and ι−_1_=0.2±0.3 s (95% CI). The low value of ι−_1_ is consistent with the time scale in which the initial increase in N_OCHRE_ takes place (Fig. 5b). The value of a_1_ corresponds to the number of photoconvertible trx-mEos2 molecules per cell within the TIRF volume and translates into 12±5 (95% CI) molecules per cell. We note that the noise term showed a much slower increase, with a characteristic time ι−_0_=8.4±0.9 s (95% CI). As a second approach, we subtracted the empirical curves N_OCHRE_-N_KO_ and obtained a difference profile (Fig. 5c) that reached a plateau at 2±1 (95% CI) fluorescent events per cell within the TIRF volume, consistent with cells transformed with ochre-trx-mEos2 expressing a minute amount of fluorescent protein copies in spite of the presence of the stop codon, and in any case larger than cells lacking the mEos2 fluorescent protein. Finally, we calculated the distributions of cumulative fluorescent events for both types of cells at time at which the plateau in the N_OCHRE_-N_KO_ difference plot had been reached (53.5 s, Fig. 5c). The distributions are clearly different (Fig. 5d). In fact, the average numbers of fluorescent events per cell within the TIRF volume (Fig. 5d) derived from the distributions are 8.1±0.5 (95% CI) (cells transformed with ochre-trx-mEos2) and 6.1±0.9 (95% CI) (knockout cells) and their difference is consistent with the a_1_ value derived from the global fittings based on equations 1 and 2.

It is important at this point to discuss the relation between the numbers of molecules detected by single-molecule localization microscopy (12±5) and the actual number of thioredoxin molecules that enable virus amplification in a cell. We detect thioredoxin molecules through the photoconversion of the attached mEos2.

Phenotypic mutations that enable the synthesis of mEos2 might on some cases replace the ochre codon at position 11 with an amino acid residue that prevents thioredoxin folding to the structure competent for interaction with viral DNA polymerase. This would obviously contribute to an overestimation of the number of thioredoxin molecules enabling virus amplification. Another source of overestimation would be related to mEos2 blinking events (Lee et al., 2012), but given the extremely low number of mEos2 molecules detected, the overestimation due to blinking cannot be but very small. In any case, the possibility of an overestimation simply reinforces the central conclusion of this work, namely that virus propagation may be enabled by an exceedingly low number of molecules. On the other hand, localization microscopy could underestimate the number of thioredoxin molecules present in the cell if some of the mEos2 proteins fail to fold properly to the photoconvertible state, thus decreasing the photoconversion efficiency. In that respect, mEos2 has been reported to have the largest photoconversion efficiency among commonly used photoconvertible and photoactivable proteins (over 60%: Durisic et al., 2014). In our hands, mEos2 displays a high folding efficiency and, indeed, UV-VIS spectra of our purified wt-trx-mEos2 fusions (Fig. 3a) indicate an approximately 1:1 fluorescent-mEos2: thioredoxin stoichiometry. Finally, it is important to note that, since successful phage T7 amplification in a cell leads to an average of hundred or more of new virions (Quimron et al., 2010), propagation could be triggered by a fraction of the cells that express numbers of thioredoxin molecules above the average value, *i.e.,* by cells at the high tail of the number of molecules distribution (see Fig. 5d). For this reason, in the following discussion we will conservatively assume that virus propagation in our experiments is enabled by a few tens of thioredoxin molecules.

Astonishing as it may seem, phage propagation enabled by a few tens of thioredoxin molecules per host cell makes sense in light of known features of the interaction of viral DNA polymerase with thioredoxin. This interaction is highly specific and very tight, likely because a highly stable complex is required to enhance processivity. In fact, the dissociation constant of 5 nanomolar has been reported for this interaction (Hamdan and Richardson, 2009). Since the volume of an *E. coli* cell is about 1.3·10^-15^ litres (see Methods), a simple calculation using the Avogadro’s number yields that 5 nanomolar is equivalent to just about 4 thioredoxin molecules per cell.

Therefore, copy numbers of a few tens of thioredoxin molecules imply cellular thioredoxin concentrations clearly above the dissociation constant for the interaction between thioredoxin and the viral DNA polymerase and, consequently, the establishment of a few tens of DNA polymerase-thioredoxin interactions (see Extended Data Fig. 6). Phage T7 has a burst size of about 100 virions and a genome size of about 4·10^4^ bp (Quimron et al., 2010). It can be easily calculated that, at an elongation rate of several hundred nucleotides per second (Tabor et al., 1987), a few tens tens of functional DNA polymerase-thioredoxin complexes could produce about a hundred copies of the phage genome in a few minutes.

Overall, this work provides a direct experimental demonstration that crucial interactions between viral and host biomolecules may need occur only a few times per host cell to enable virus propagation. Rare but crucial virus-host biomolecular interactions are not of course limited to the recruitment of host factors. For instance, virus entry in a host cell is commonly triggered by a single interaction event involving a viral protein and the adequate receptor on the surface of the host cell. Also, viruses may need to block host antiviral factors (Kluge et al., 2015), which in some cases could be present at low copy numbers.

An immediate implication of the capability to use biomolecules at very low copy numbers is that viruses may readily take advantage of pre-existing low-level protein diversity. Errors during gene expression provide a major source of such protein diversity. Transcription and translation errors are common and lead to the so-called phenotypic mutations (Drummond and Wilke, 2009; Goldsmith and Tawfik, 2009; Evans et al., 2018; Landerer et al., 2024) which are orders of magnitude more common than genetic mutations linked to changes in DNA sequences. The diversity of protein variants generated by gene-expression errors can be illustrated with a simple back-of-the-envelope calculation. Copy numbers for human proteins range between ∼10^2^ and ∼10^7^, with the maximum of the distribution at about 10^5^ molecules per cell (Kulak et al., 2014). Using a recent estimate of about 20% for the fraction of protein molecules bearing phenotypic mutations (Landerer et al., 2024), it is easily calculated that, out of 10^5^ copies per cell of a given representative protein, around 20000 would be variants harbouring phenotypic mutations. For comparison, this number of variants is about 3 times the total number of possible single amino acid replacements (19×375) in a human protein of median length (Brocchieri and Karlin, 2005).

Certainly, phenotypic mutations are not inherited. Therefore, they are neither purged nor amplified by natural selection. This means that the diversity of variants continuously generated by gene-expression errors is not pruned by selection and remains available as a resource for adaptation. Viruses may readily exploit this “hidden” resource because, as shown here, they can amplify in a host cell on the basis of a few successful biomolecular interactions enabled by protein molecules present at exceedingly low levels. Moreover, virus amplification does not need to occur in all, or even most, host cells to sustain propagation. This is because virus amplification in a single host cell can lead to many new virions, as shown by reported burst sizes that range from several tens to several thousands of virions, depending on the virus and the host studied (Phillips and Milo, 2015). Overall, a few successful biomolecular interaction events in just a few host cells may lead to the generation of many new virions thus triggering propagation. Crucial interactions could be mediated in some cases by host or viral protein variants resulting from gene expression errors and present at exceedingly low levels. Our results do support this scenario, as we observe virus propagation linked to a few host factor molecules which, given our experimental design, are necessarily generated through mistranscription or mistranslation. Certainly, virus proliferation based on protein variants present at exceedingly low levels may not be efficient. Crucially, however, this inefficient propagation may make it possible for the virus to meet new challenges by allowing its survival until mutations that provide inheritable adaptation appear at the genetic level. This “look ahead” evolutionary role for phenotypic mutations was in fact proposed years ago on theoretical grounds (Whitehead et al., 2008).

The notions expounded above are conveniently illustrated with the phenomenon of cross-species transmission. Viruses occasionally jump between species, sometimes with disastrous consequences for the new hosts. Cross-species transmission is puzzling from a molecular point of view because the “jumping virus” needs to establish effective biomolecular interactions simultaneously in the old and the new hosts. However, the biomolecules involved (cell surface receptors, host factors to be recruited, antiviral molecules to be blocked) may substantially differ between the two hosts in terms of amino acid sequence and, therefore, in terms of the chemical composition of the exposed molecular surfaces available for biomolecular interaction. Certainly, suitable mutations will lead to adaptation to the new host. Yet, virus-host biomolecular interactions are extensive (Watanabe et al., 2014; Ramage and Cherry, 2015; Baggen et al., 2021;) and evolution has no foresight (Jacob, 1977). That is, the, presumably large, number of adaptive mutations required will be fixed only after the virus has “jumped”, it is already replicating in the new host and the mutations confer a selective advantage. The question is, how does the virus start to replicate in the new host in the first place, *i.e.*, before the adaptive mutations have appeared at the genetic level? Phenotypic mutations may provide a way out of this catch-22. Errors during gene expression generate a large diversity of variants of host and viral proteins bearing a wide variety of phenotypic mutations. This may well provide a “hidden” resource that viruses can exploit to initially establish the required biomolecular interactions in the new host, thus allowing survival and giving natural selection a chance to act.

The plausibility of the scenario proposed in the previous paragraph can be illustrated with some simple order-of-magnitude calculations on SARS-CoV-2. There are about 300 copies of the spike monomer on the surface of a SARS-CoV-2 virion and infection of a host cell is likely triggered by an interaction of a spike molecule with an ACE2 receptor at the cell surface (Bar-On et al., 2020). Cross-species transmission requires that the virus spike can interact with the ACE2 receptor of the new host.

Mutations in the spike linked to cross-species transmission have been identified (Zech et al., 2021; Tan et al., 2022). The question is, how does the virus “sets foot” in the new host before such mutations have appeared at the genetic level? The infective dose of SARS-CoV-2 has been estimated on the order of hundreds of virions for normal transmission in humans (Karimzadeh et al., 2021) but most COVID-19 animal infection models, which may be more representative of cross-species transmission in this context, require doses in the 10^4^-10^6^ range (Brosseau et al., 2022). 10^4^-10^6^ virions could carry up to several hundred million copies of the spike monomer, out of which, using the 20% estimate (Landerer et al., 2024), many millions would be variants with phenotypic mutations. It is plausible, therefore, that a fraction of these variants harbour phenotypic mutations that enable the interactions with the ACE2 of the new host. Of course, the enabling interactions could also be mediated by phenotypic mutations in ACE2. Using for illustration a reported value for the copy number of ACE2 in HeLa cells of ∼30000 (Kulak et al., 2014), about 6000 ACE2 molecules per cell would harbour phenotypic mutations. Overall, it appears plausible that phenotypic mutations on either side of the crucial spike-ACE2 interaction enable the infection of a significant number of host cells. This could suffice to trigger propagation since each of the new-host infected cells would yield an average of 1000 new virions (Bar-On et al., 2020). It is relevant here that ∼1000 new virions mean about 300000 new copies of the spike monomer, out of which, using again for illustration the 20% estimate (Landerer et al., 2024), around 60000 would be variants with phenotypic mutations.

Similar order-of-magnitude calculations support the plausibility of a role of phenotypic mutations in the surprising capability of viruses to evade antiviral strategies, as most clearly exemplified by the so-called “influenza puzzle” (Ellebedy and Ahmed, 2016), *i.e.,* the fact that influenza remains a health problem despite repeated exposure of the population to viral proteins through natural infection and vaccination. There are about 1500 copies of the hemagglutinin monomer on the surface of an influenza virion and infection of a host cell is likely triggered by an interaction of the globular head of the hemagglutinin molecule with sialic acid at the cell surface (Sautto et al., 2018). Some antibodies bind to the globular head and block interaction with sialic acid thus preventing infection, but single mutations in hemagglutinin that evade antibodies are known (Doud et al., 2018). Some studies have reported a minimal infective dose for influenza virus of about 1000 virions (Nikitin et al., 2014). Using this number for illustration, 10^3^ virions could carry more than a million copies of the hemagglutinin monomer out of which, using the 20% estimate (Landerer et al., 2024), a few hundred thousand would be variants with phenotypic mutations. It appears highly plausible that some of these variants evade antibody binding and enable infection. Infection of just a few cells could trigger propagation, since burst sizes for influenza virus are reported between 500 and 10000 (Phillips and Milo, 2015).

Therefore, infection of just a single cell could generate many millions of copies of the hemagglutinin monomer, out of which many hundred thousand would be variants with phenotypic mutations. Certainly, these are back-of-the-envelope calculations that can be modified in several ways, including for instance that, in real transmission scenarios, the infective doses could much higher than the minimal one. The general point is that viruses can achieve huge amplifications on the basis of a few biomolecular interactions, which makes them particularly suited to take advantage of the protein diversity arising from gene-expression errors.

It remains now to make some suggestions as to how the role of phenotypic mutations on virus adaptation could be experimentally addressed. One interesting possibility in this context is that phenotypic mutations caused by transcription errors are particularly relevant. It must be noted first that difference between genetic mutations and phenotypic mutations linked to transcription errors applies even to RNA viruses, since the viral RNA-polymerase would generate both genomic RNA to be encapsulated in the virions and mRNAs to be used in protein synthesis. Certainly, the transcription error rate is low as compared with the translation error rate (Drummond and Wilke, 2009). However, transcription errors may have a stronger impact at the protein level because each mRNA molecule is typically translated many times (Traverse and Ochman, 2016). Next-generation sequencing methodologies with the capability to determine mutations present in RNA at very low level are available (Lu et al., 2020).

Furthermore, transcription errors are known to be far from completely random (Gout et al., 2017; Acevedo et al., 2014). It would seem then feasible to determine the landscape of transcription errors for viral (and host) proteins and to assess the extent to which the most prevalent phenotypic mutations enable biomolecular interactions crucial for virus propagation.

## Methods

### Cells

*E. coli* expresses thioredoxin 1 (which we refer to in the main text simply as “thioredoxin”) and thioredoxin 2, which is induced under oxidate stress and includes a zinc-binding domain (Delgado et al., 2017). To our knowledge, thioredoxin 2 does not act as a processivity factor for the viral DNA polymerase. In any case, the knockout Trx^-^ strain used in this work (originally a gix from John Beckwith, Harvard Medical School) is deficient in both thioredoxin genes. Genes encoding thioredoxin and thioredoxin-fluorophore constructs (both with and without stop codons at position 11in the amino acid sequence) were synthesized with a His-tag at the C-terminal and cloned into pET30a(+) plasmid with kanamycin resistance (GenScript Biotech). Transformation of knockout Trx^-^ cells and other details are essentially as we have previously described (Delgado et al., 2017).

Purification of the trx-mEos2 construct for determination of its UV-VIS spectrum (Fig. 3a) was carried by Ni-NTA affinity chromatography, followed by passage through PD10 columns to obtain protein solutions in Hepes 50 mM, pH 7. Protease inhibitors were added in all steps of the purifica{on. Three independent purifica{ons were carried out, leading to an average value of 1.45±0.30 for the 506 nm vs. 280 nm absorbance ratio. The absorbance peak at 506 nm is due to the fluorophore and has a reported ex{nc{on coeffcient of 5.6·10^4^ M^-1^cm^-1^ (McKinney et al., 2009). The absorbance peak at 280 nm reflects protein contributions from both mEos2 and thioredoxin. Published spectra indicate ex{nc{on coeffcients at 280 nm of about 2.5·10^4^ M^-1^cm^-1^ for mEos2 (McKinney et al., 2009) and of 1.4·10^4^ M^-1^cm^-1^ for thioredoxin (Georgescu et al., 2001). The fusion trx-mEos2 construct, therefore, is expected to have an ex{nc{on coeffcient of 3.9·10^4^ M^-1^cm^-1^ at 280 nm and show a 506 nm vs. 280 nm ratio of 1.44 in agreement with our experimental value of 1.45±0.30.

### Plaque assays

Plaque assays were carried out as previously described in Luzon-Hidalgo, et al. (2021).

### Lysis experiments

Lysis experiments were performed using a simplified adaptation of a previously employed protocol (Luzon-Hidalgo et al., 2021). Briefly, overnight preinocula were incubated in LB medium at 37 °C, which, for transformed cells, had been supplemented with kanamycin (50 mg/mL). Cultures were diluted 1/200 in fresh medium and, when the absorbance at 600 nm had reached 0.25-0.30, microliter volumes of virus stock solution were added to the desired Multiplicity of Infection (MOI). Cultures were incubated for 1 hour at 37 °C and absorbance was measured again. In experiments with very high MOI values, the absorbance values at 600 nm measured after 1 hour were very low, on the order of 10^-2^-10^-3^, indicating that essentially all cells had been killed. The survival cell fraction at each given MOI was calculated from the absorbances at 600 nm using the following equation:

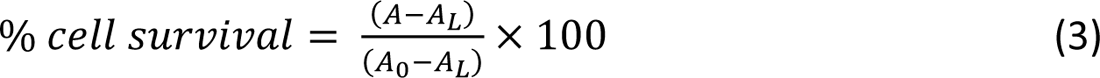

where A_0_ and A are, respectively, the initial absorbance and the absorbance at 1 hour in one experiment corresponding to a given MOI value, and A_L_ is the (very low) absorbance at 1 hour in another experiment in which a very high MOI value was used and essentially complete cell death was attained after 1 hour. MOI values were calculated from the concentration in the virus stock solution in plaque-forming-units (PFU) per unit volume. This concentration was calculated for each amplified phage using plaque assays with a DHB3DE3 *E. coli s*train as previously described (Delgado et al., 2017).

### Single molecule experiments

Bacterial cultures were grown for microscopy experiments on a sterilized chambered 1.5H glass bottom coverslip with 8 individual wells coated with poly-L-lysine. 350 µL of a 1/1000 dilution from an over day preinocula, that had reached exponential growth phase, were transferred to each well. These cultures were incubated overnight and protected from light at room temperature. The next day, the wells were washed to eliminate floating and non-immobilized bacteria before fixation with 4% paraformaldehyde and sodium borohydride (NaBH_4_) to decrease autofluorescence.

Finally, 250 µL of a 1/1000 dilution in PBS of 80 nm gold colloid beads was added to the wells that allowed image drift correction when necessary. Experiments were carried out on a TIRF microscope (Leica Microsystem), with an oil-immersion objective HC PL APO 160x 1.43 NA, an Orca Flash 4.0 sCMOS camera (Hamamatsu), a Gemini W-View splitter (Hamamatsu), and high-power lasers suitable for single molecule localization microscopy techniques. An additional high pass filter (OG530) was located at the microscope condenser to prevent spontaneous mEos2 photoconversion during sample manipulation and focusing.

Single images of mEos2 before photoconversion were acquired at 488 nm at minimal to prevent mEos2 copies from bleaching. mEos2 was photoconverted by continuously illuminating the sample at 405 nm throughout the length of the experiment using either 1 or 0.1 mW/mm^2^ power density depending on the sample. Samples were simultaneously imaged at 561 nm (20 W/mm^2^) using a multi-band bandpass filter (excitation: 397-413 nm; 481-495 nm; 550-566 nm; 627-643 nm. Emission: 420-480nm; 500-550 nm-580-630 nm; 660-860 nm) at the microscope and extra bandpass filters at the GEMIN image splitter (Semrock FF01-512/25-25; FF01-630/92-25). The exposure time was 100 ms. All images were acquired on TIRF configuration (150 nm penetration depth) with 102-nm pixel at the sample plane (256×256 pixels). The total length of the imaging sequence was either 5 min or 10 minutes (depending on the sample).

### Confocal microscopy

Bacterial cultures were grown as single molecule experiments and living cells, without sample fixation nor quenching, were analysed by confocal microscopy. Images were acquired using a Leica STELLARIS 5 multispectral confocal system (Leica Microsystems) with an oil immersion HC PL APO 63x 1.40 NA objective (Leica Microsystem). Samples were excited at 488 nm using either 4 W/mm^2^ or 20 W/mm^2^ at the sample plane depending on the sample. Fluorescence emission was collected using a hybrid (HyD) S detector in the range of 498 nm-620 nm, gain at 2.5% or 122.79% depending on the sample, and 52 nm pixel.

### FRAP experiments

Bacterial cultures were grown and prepared for analysis as previously described for single molecule experiments. For the current experiments, no incubation with 80 nm gold colloid beads has been performed.

Fluorescence recovery after photo bleaching (FRAP) experiments were performed using a Leica STELLARIS 8 STED 3X multispectral confocal nano and mesoscopy system (Leica Microsystems) using a water immersion HC PL APO CS2 63x 1.20 NA 63x objective. FRAP imaging was performed using the FRAP AB mode of the Leica LAS X software. Pre-bleaching and post-bleaching imaging was performed at 491 nm and fluorescence emission was collected using a hybrid spectral (HyD S1) detector in the range of 499 nm-594 nm. Bleaching was achieved at 491 nm with a 30 W/mm^2^ laser power density, using zoom in mode. For each cell a 0.07 µm^2^ bleaching area was defined, and the FRAP protocol was established as following: pre-bleaching one-image acquisition, followed by 15 seconds bleaching and an 80-second post-bleaching recovery phase.

### Single-molecule data analysis

Molecules were localised using Thunderstorm (Ovesný, et al. (2014)) plugin for ImageJ. Camera parameters were specifically configured with a pixel size of 101.6 nm, 0.46 photoelectrons per A/D count, and a base level of 1650 ADU. Fitting parameters used were difference-of-Gaussians filter, local maximum detection of single molecules and integrated Gaussian weighted least squares fitting. Post-processing steps include drift correction by fiducial markers and merging frames (unlimited).

Cells were automatically identified and segmented based on the phase-contrast and 488-excitation (before photoconversion) images using a deep-learning 2D-segmentation algorithm (Misic, Panigrahi et al. 2021). We visually supervised the segmented cells, amended the cell contours were needed and interfaced them with Fiji (Schindelin et al. 2012) using the Roifile Python Library (Gohlke, 2024). We then assigned every localization in the PALM image sequence to its corresponding cell (ROI) and built the empirical cumulative sum of fluorescent events per cell.

Finally, we fittted a cumulative (double/single) exponential - linear model (equations 1 and 2 in the main text) to the identifications cumulative sum, weighting the optimization with the uncertainty of each point. To estimate the uncertainty of the fitted parameters (a_0_, a_1_, ι−_0_, ι−_1_, b), we randomly resampled seven times (without replacement) the ochre and knockout data sets (575 and 193 cells, respectively). This way, we produced seven subsets of approximately 82 (ochre) and 27 (knockout) cells each. We then global-fitted equations 1 and 2 to the subsets as explained above. We finally computed the uncertainty of the fitted parameters as the standard error of the mean of the obtained seven values and multiplied this value timers 1.96 to derive the 95% confidence interval. This resampling procedure was repeated at least five times with identical results within the experimental uncertainty. Image analysis and fitting algorithms developed for this study are available at https://github.com/cnbbiophot.

### Determination of the cell volume

We measured the long (*L*=2.02±0.04 µm) and short (*w*=0.96±0.01 µm) cell dimensions from the phase contrast images using the Fit Ellipse tool in Fiji. We measured 510 cells and assumed sphero-cylinder morphology (eq. 4) to compute the cell volume, resulting in *V=*1.30±0.06 fl. All errors are 95% CI.

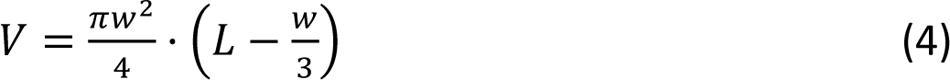

## Acknowledgements

This research was supported by Grant PID2021-124534OB-100 (to J.M. S.-R.) funded by MICIU/AEI/10.13039/501100011033, Grant IHRC22/00004 (to J.M.S.-R.) funded by the “Instituto de Salud Carlos III (ISCIII)” and Next Generation EU, Grant PID2021-125024NB-C21 funded by MCIN/AEI / 10.13039/501100011033 / FEDER, UE awarded to JRI, SEV-2017-0712 funded by MCIN/AEI/10.13039/501100011033 and Grant PRE2019-089850 (to R.L.-H.) funded by MICIU/AEI/10.13039/501100011033 and by “ESF Investing in your future”

The authors declare no competing interests

---

**Extended Data Fig. 1.**
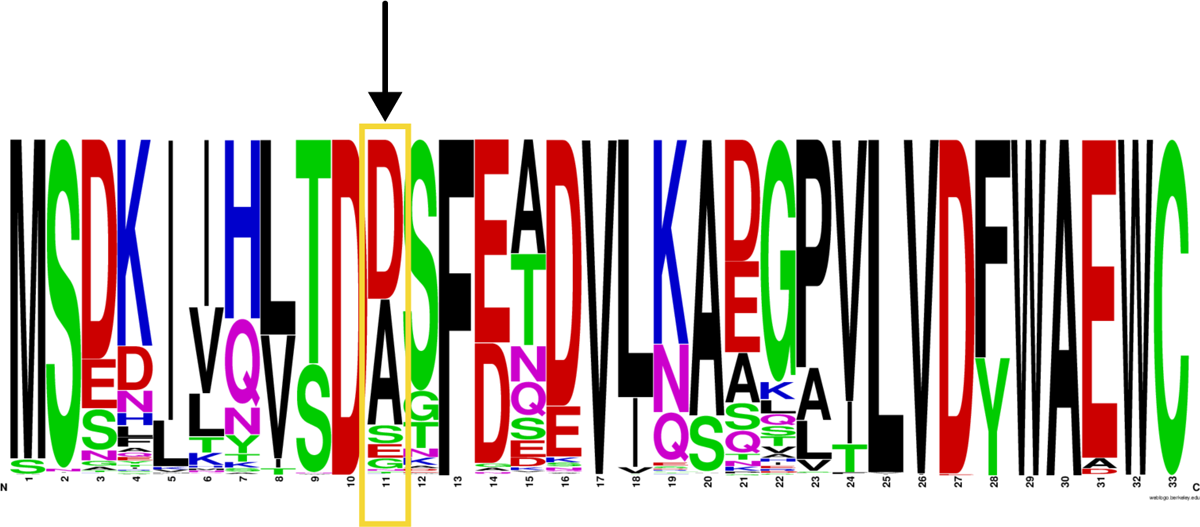
Sequence logo for thioredoxin. A blast search was performed using the sequence of *E. coli* thioredoxin was performed. The sequences with identity with the query higher than 70 % were retained for residue frequency calculation. Position 11 is exposed to the solvent in the 3D-structure of thioredoxin (Fig. 1b). An aspartate is present at position 11 in *E. coli* thioredoxin. However, the frequency analysis summarized in the sequence logo indicates that position 11 in thioredoxins can admit other amino acid residues. Position 11 was selected for stop-codon insertion.

**Extended Data Fig. 2.**
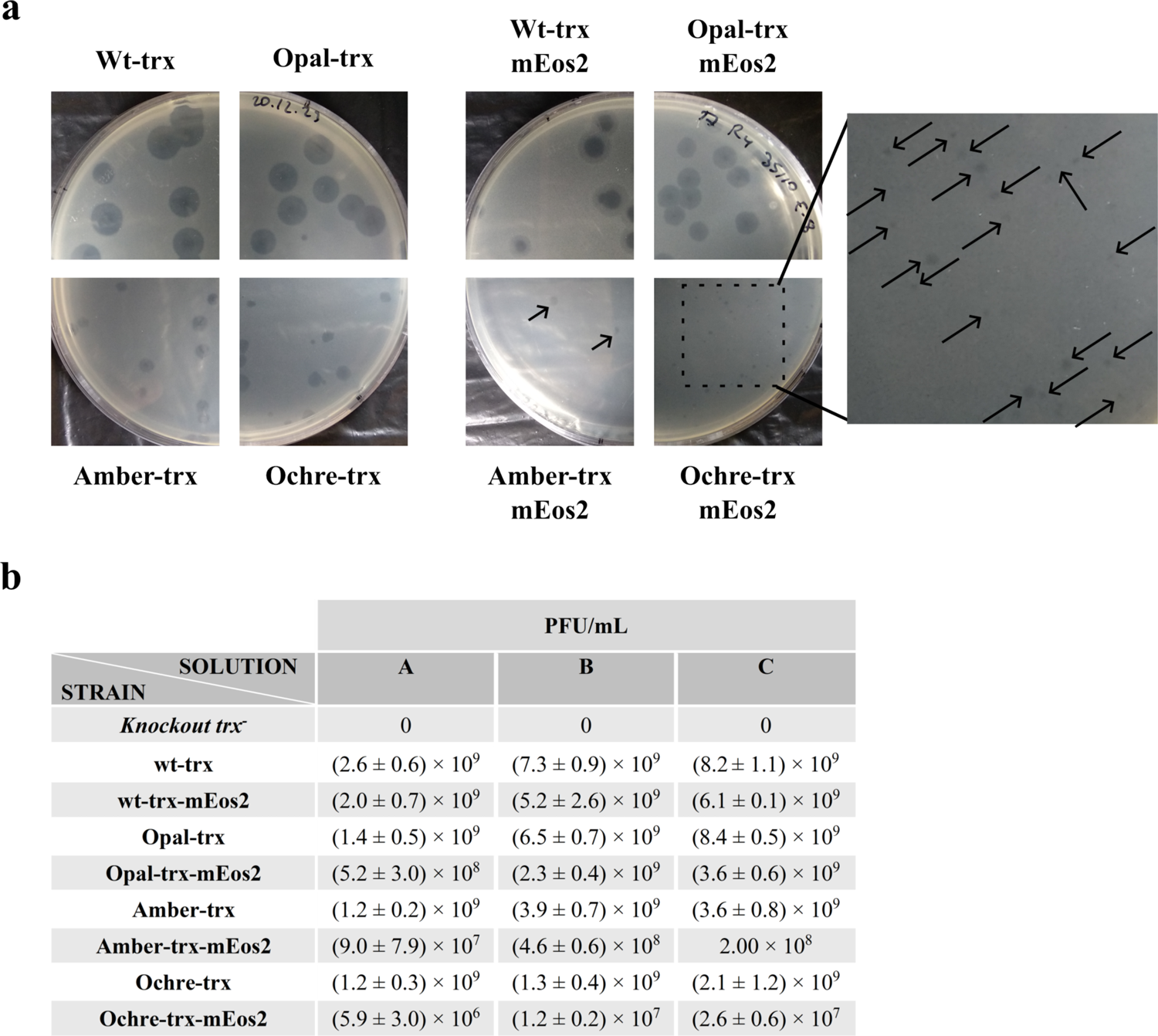
Determination of plaque-forming units (PFU) for phage solutions. **a**, representative examples of T7 plaque formation after inoculating various hosts with the phage. The labels indicate the gene with which the knockout Trx^-^ had been transformed. Plaques observed with cell transformed with ochre-trx-mEos2 are small and blowup is provided. **b,** number of PFUs per millilitre for 3 phage T7 solutions (labelled A, B and C) derived from 3 independent amplifications. Various strains were inoculated with the virus solutions. Except for the knockout Trx^-^, the strains are identified by the gene used in the transformation. Values of PFU/mL for the phage solutions were calculated from the numbers of plaques upon serial dilutions, as described in Luzon-Hidalgo et al. (2021). Average values and standard deviations from three determinations are given.

**Extended Data Fig. 3.**
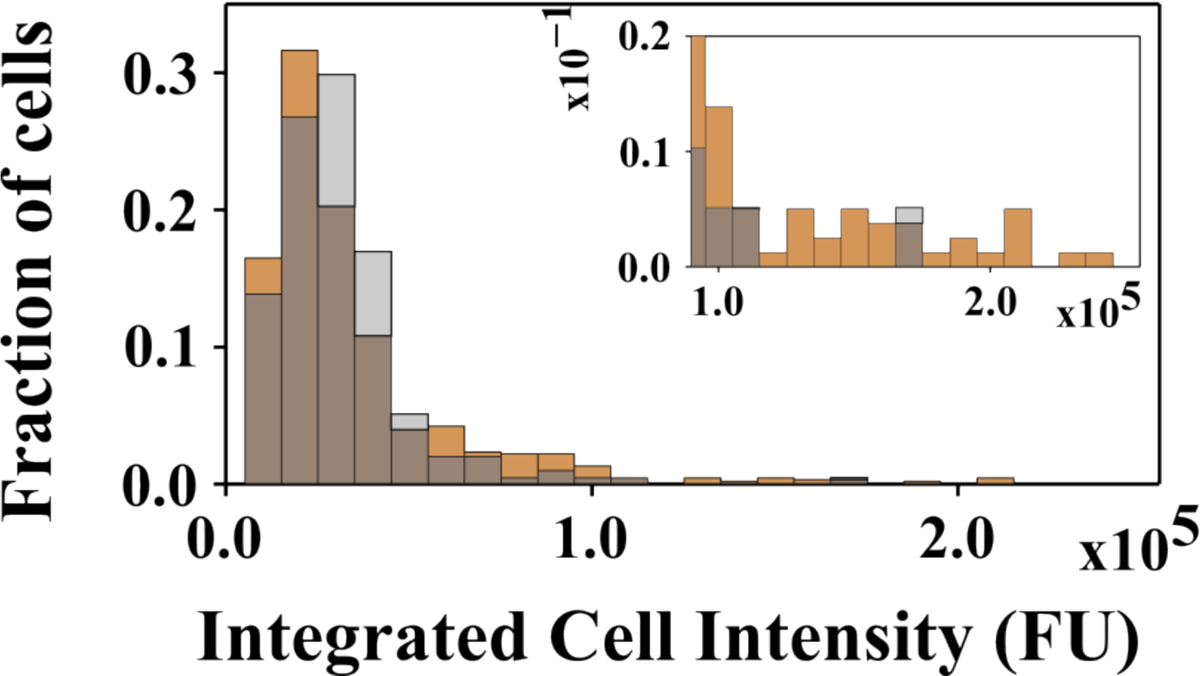
Distribution of fluorescence intensity for cells transformed with ochre-trx-eGFP. The data (ochre) are shown superimposed with those corresponding to the knockout Trx^-^ cells (grey). The distributions show that the fluorescence levels for upon transformation with ochre-trx-eGFP are similar, for almost all cells, to those observed with the knockout *E. coli* Trx^-^ cells, indicating that the contributions from endogenous autofluorescence and eGFP are comparable. Yet, the blowup of the high fluorescence intensity region (inset) shows a prevalence of ochre-trx-eGFP transformed cells and supports that a small fraction of these cells expressed eGFP to a detectable level in these experiments. Overall, however, it does not appear possible to use eGFP fluorescence to arrive at an estimate of the average number of molecules generated per cell.

**Extended Data Fig. 4.**
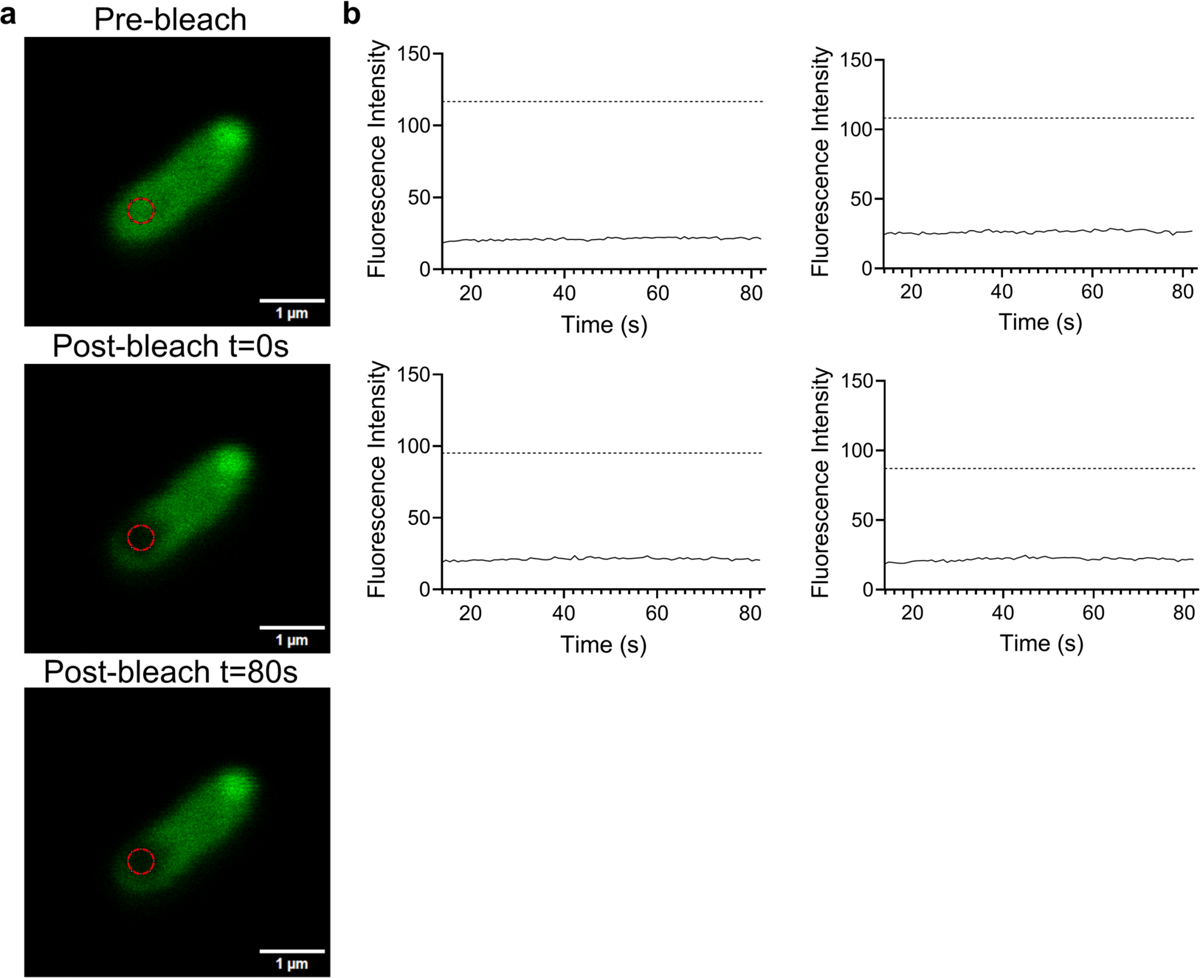
FRAP experiments in bacteria expressing wt-trx-eGFP. **a**, Representative confocal images of an eGFP-expressing bacterium fixed following the same protocol as used for single-molecule experiments. Top panel: the bacterium before bleaching; middle panel: immediately after the beaching cycle; bottom panel: 80 seconds after bleaching. The bleaching area is highlighted in red. **b,** Representative FRAP curves of four different bacteria. Solid lines are the eGFP mean fluorescence intensity within the bleached area over 80 seconds. Dotted lines describe the eGFP mean fluorescence intensity of the pre-bleached area. There is no significant fluorescence recovery after 80 seconds, indicating that protein diffusion is negligible.

**Extended Data Fig. 5.**
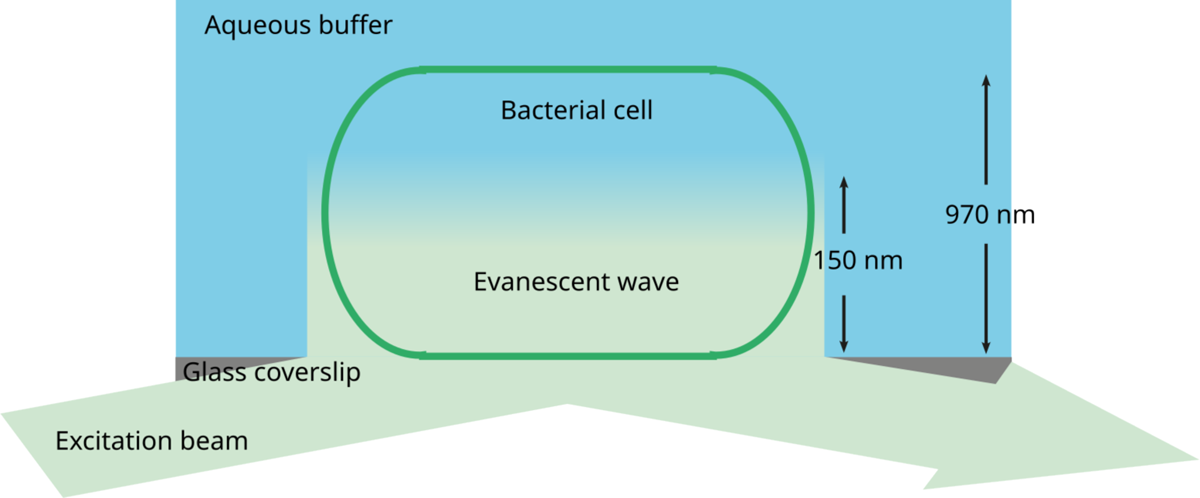
Total Internal Reflection Microscopy (TIRF). The excitation beam arrives at the coverslip-aqueous buffer and is totally reflected into the buffer if the incidence angle is larger than the critical angle. In this case, an evanescent wave travels along the interface with an amplitude that falls off exponentially with distance from the interface. Consequently, fluorescent molecules are only efficiently excited in the region closest to the interface (≈150 nm on our microscope). TIRF is, thus, particularly suited for single-molecule fluorescence studies, since background fluorescence is extremely low. The breadth of the bacteria used in this work is ≈970 nm high implying that fluorescent molecules in the bacteria cytoplasm but outside the TIRF excitation volume will not get excited. For this reason, the count of fluorescent molecules within the TIRF volume must be scaled by a factor 6.5 to derive the total molecule count inside the bacterium. Note that an illustrative scheme is displayed here and that the lengths are not shown to scale.

**Extended Data Fig. 6.**
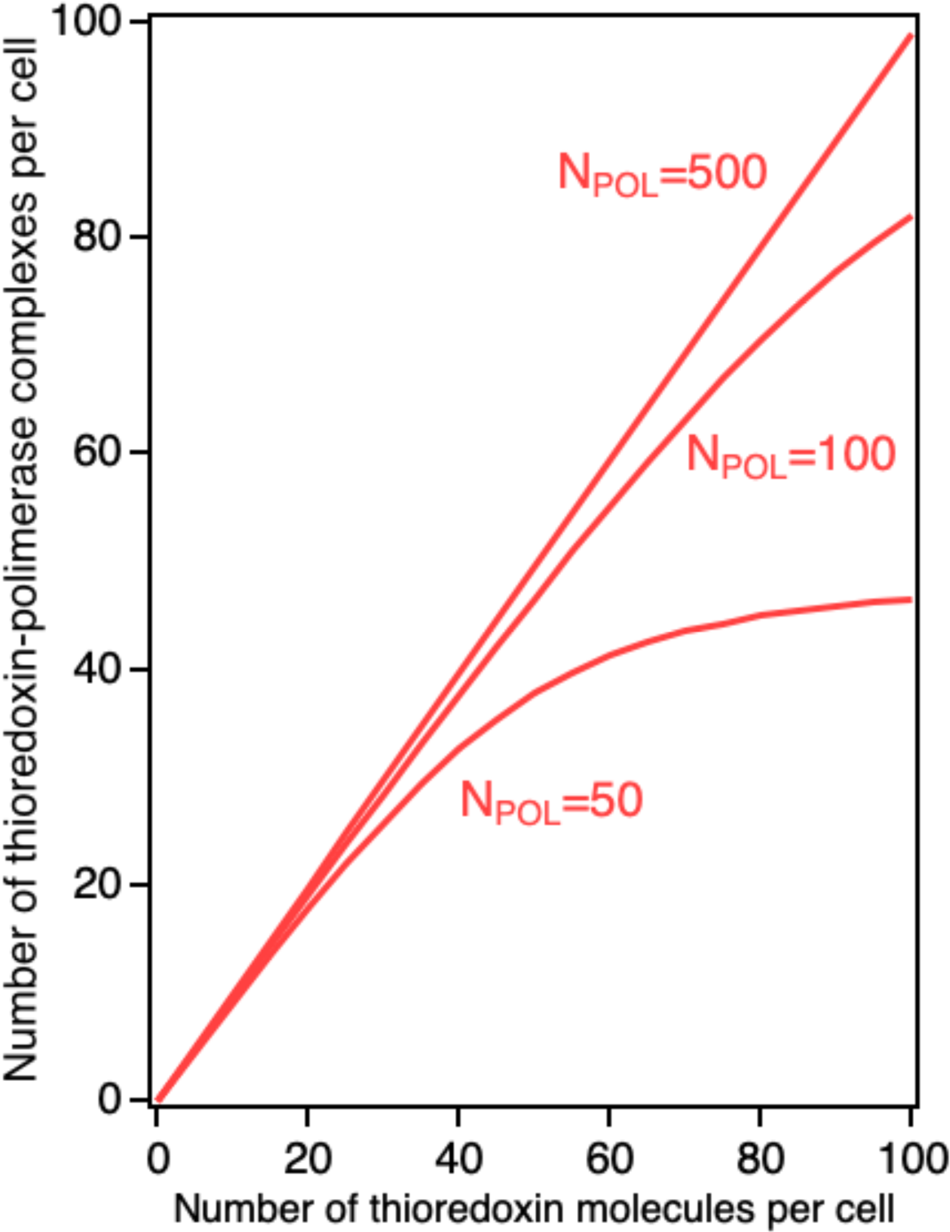
Thermodynamic estimation of the number of thioredoxin-viral DNA polymerase complexes within a cell. The interaction between the two proteins to yield the complex, pol+trxΗpol-trx is described by a dissociation constant, K_D_=[pol][trx]/[pol-trx], of 5 nanomolar (Hamdan and Richardson, 2009). The concentrations of free (non-bound) polymerase and thioredoxin, [pol] and [trx], that appear in the equation for K_D_ are related to the corresponding total concentrations through mass balance: [pol]_T_=[pol]+[pol-trx] and [trx]_T_=[trx]+[pol-trx]. These two equations can be solved for [pol] and [trx], and subsequent substitution into the expression for K_D_ yields a second-order equation that can be solved for [pol-trx] as a function of [pol]_T_ and [trx]_T_. Finally, concentrations can be converted to numbers of molecules using Avogadro’s number and the cell volume. We have used this procedure to calculate profiles of number of complexes versus number of thioredoxin molecules within a single cell for several values of the number of viral DNA polymerase molecules in the cell (N_POL_). It is likely that, upon infection of a cell, the copy number of viral DNA polymerase reaches very high values. Yet, we have purposedly used low N_POL_ values for these calculations. The profiles obtained show that a dissociation constant of 5 nanomolar predicts that, even with low N_POL_ values, a few tens of thioredoxin molecules in a cell should lead to a few tens of thioredoxin-viral DNA polymerase complexes in the cell.

## Notes

### Competing Interest Statement

The authors have declared no competing interest.

